# Tanycyte-mediated synapse pruning shapes hypothalamic circuits controlling metabolism

**DOI:** 10.1101/2025.11.14.688419

**Authors:** Rafik Dali, Irina Kolotueva, David Lopez-Rodriguez, Chaitanya K. Gavini, Tamara Deglise, Fanny Langlet

## Abstract

Synaptic pruning by glial cells is critical for refining neural circuits, yet which glial populations mediate this process in hypothalamic networks controlling energy balance remains poorly understood. Here, using 3D-ultrastructural analyses and molecular approaches, we show that tanycytes —specialized glial cells lining the third ventricle— actively engulf excitatory synapses in the mediobasal hypothalamus during a defined postnatal window. This phagocytic activity relies on tanycytic MFG-E8 and phosphatidylserine exposure on synaptic elements. Disruption of this pathway reduces synapse uptake *in vitro* and alters body weight control around weaning and glucose homeostasis in adulthood.

These findings identify tanycytes as previously unrecognized glial mediators of synaptic pruning in hypothalamic circuits, establishing their active role in shaping neural networks underlying energy balance.

## INTRODUCTION

The hypothalamus is a central hub that integrates diverse sensory, metabolic, and hormonal inputs to regulate essential physiological functions, including energy balance, thermoregulation, reproduction, stress responses, and circadian rhythms^1^. Proper regulation of these functions relies on precisely wired neuronal circuits^1,2^, which are established through sophisticated developmental processes that begin during embryogenesis and continue into postnatal life^3,4^. This process requires a tightly orchestrated sequence of cellular and molecular events^3,5^, including the generation of newborn neurons^6,7^, the specification of neuronal identities^7^, the establishment of long-range axonal projection^3,8^, and the formation, stabilization, and elimination of synaptic connections^9,10^. While the early stages of hypothalamic development – including neurogenesis, neuronal migration, and axonal projection – have been increasingly characterized^3,8^, the cellular and molecular mechanisms governing the establishment of precise and functionally efficient synaptic connectivity remain largely unresolved.

One critical step in postnatal circuit maturation is synaptic pruning, a neuronal activity-dependent process in which excess or weak synapses are selectively eliminated^11^. In well-studied regions such as the visual cortex and hippocampus, pruning is mediated by glial cells, primarily microglia^12,13^ and, to a lesser extent, astrocytes^14^. These cells engulf pre- and postsynaptic elements through receptor-mediated recognition of "eat-me" signals, such as phosphatidylserine^15^, often aided by bridging molecules like MFG-E8 ^16,17^. This glia-mediated synapse elimination not only shapes developing circuits in response to neuronal activity^13^ and metabolic cues^18^, but also occurs in the adult brain in response to environmental stimuli^19^. Despite this growing understanding, the mechanisms governing synaptic pruning in the hypothalamus – a brain region with distinctive cytoarchitecture, lifelong plasticity, and crucial homeostasis roles^20–22^ – remain largely undefined.

Among hypothalamic glia, tanycytes are uniquely positioned to influence circuit maturation^23^. Tanycytes are specialized ependymal cells lining the wall of the third ventricle and extending long processes deep into hypothalamic nuclei^23,24^. While traditionally associated with blood-brain exchange regulation^25,26^, metabolic sensing^27–30^, and neuronal function modulation^27,31^, anatomical and ultrastructural studies have also revealed that tanycytic processes are in close proximity to synapses in the mature brain^32,33^. Notably, tanycytic processes can completely envelop presynaptic terminals within a double-membrane vesicle, raising the possibility that these cells could directly contribute to synaptic remodeling^32^. Developmentally, tanycytes arise during late embryogenesis, starting at embryonic day 12 in mice, and continue to be generated in a gradual and regionally heterogeneous manner into the early postnatal period^34,35^. Then, their maturation takes place over the first two postnatal weeks, reaching completion by approximately postnatal day 14 (P14) in mice^34,35^. This period of maturation coincides with a key window of synapse formation and refinement in the mediobasal hypothalamus^3^, suggesting that tanycytes could participate in shaping local circuits during a critical developmental phase.

Despite their strategic location, morphology, cytoarchitecture, and developmental timing, the role of tanycytes in synapse remodeling has remained unexplored. Here, we demonstrate that tanycytes act as phagocytic cells during a defined developmental window, engulfing excitatory synapses through a mechanism requiring the expression of the bridging molecule MFG-E8 and the exposure of phosphatidylserine on synaptic elements. Disruption of this pathway reduces synapse uptake *in vitro* and leads to physiological alterations around the time of weaning and in adulthood *in vivo*. Our findings position tanycytes as novel glial contributors to synaptic refinement and establish their role in shaping hypothalamic circuits essential for homeostatic regulation.

## RESULTS

### Tanycytes actively engulf excitatory synapses during postnatal hypothalamic development

To define the timing of synapse development and refinement in the arcuate nucleus of the hypothalamus (ARH), we first quantified the density of glutamatergic and GABAergic synaptic terminals— including pre-synaptic (VGLUT2⁺ or VGAT⁺), post-synaptic (PSD95⁺ or Gephyrin⁺), and colocalized (VGLUT2⁺/PSD95⁺ or VGAT⁺/Gephyrin⁺) elements— at several postnatal time points (Fig. 1a-h). Regarding glutamatergic synapses (Fig. 1a-d), the number of colocalized elements remains stable from postnatal day 4 (P4) until P13, followed by a marked decrease at P14–P15 (Fig. 1d), suggesting active synapse elimination during this developmental window. The number of pre-synaptic elements exhibits a similar temporal profile (Fig. 1b), whereas the number of post-synaptic elements remains stable (Fig. 1c). In contrast, GABAergic synapse number shows a modest increase from P4 up to P13 (Fig. 1e-h), with only a slight decrease at P15 (Fig. 1h). Presynaptic element density is stable from P4 to P13 before decreasing at P15 (Fig. 1f), while the postsynaptic element density increases at P14–P15 (Fig. 1g). As described previously^9,10^, hypothalamic circuit formation and refinement during the first two postnatal weeks in mice primarily involves glutamatergic synapses, coinciding with maturation of excitatory circuits.

**Figure 1.**
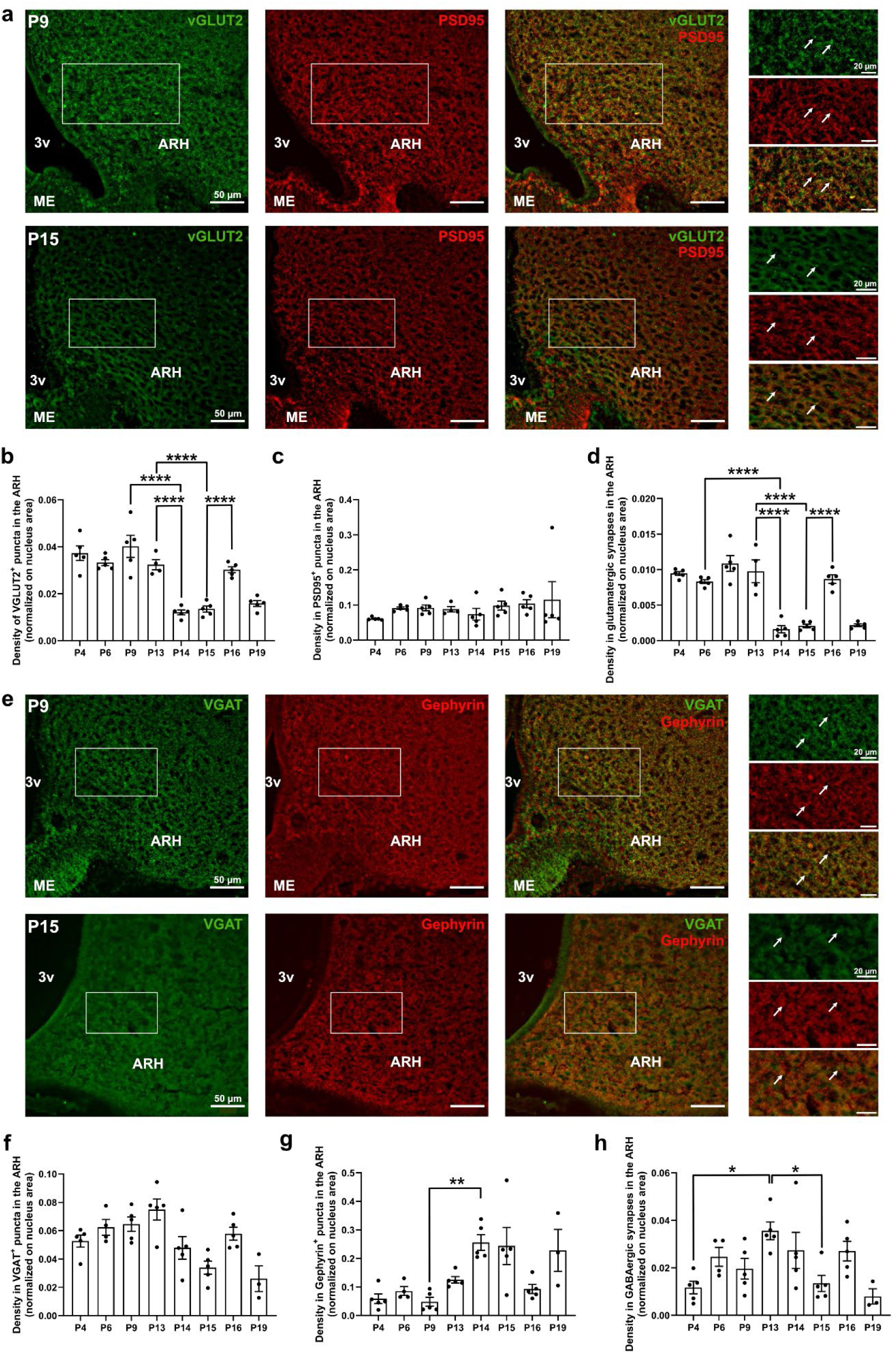
Synaptic density in the ARH is temporally regulated during postnatal development. **a**, Representative confocal images showing VGLUT2^+^ (green), PSD95^+^ (red), and merge puncta (yellow) in the arcuate nucleus (ARH) at postnatal days 9 (P9) and 15 (P15). **b–d**, Quantification of VGLUT2^+^ puncta (b), PSD95^+^ puncta (c), and putative synapses (colocalized VGLUT2/PSD95; d) in the ARH across postnatal development. **e**, Representative confocal image showing VGAT^+^ (green), Gephyrin^+^ (red), and merge puncta (yellow) in the arcuate nucleus (ARH) at postnatal days 9 (P9) and 15 (P15). **f–h**, Quantification of VGAT^+^ puncta (f), Gephyrin^+^ puncta (g), and putative synapses (colocalized VGAT/Gephyrin; h) in the ARH across postnatal development. 3v, third ventricle; ARH, arcuate nucleus; ME, median eminence; Scale bars: as indicated. Data represent mean ± s.e.m.; n = biological replicates (black dots in b-d and f-h) ; statistical details in Methods. *, p<0.05; **, p<0.01; ***, p<0.001; ****, p<0.0001.

To examine whether tanycytes participate in this process, we induced tdTomato expression in neonatal pups to visualize the entire morphology of individual tanycytes (Fig. 2a) and assess their interactions with synapses (Fig. 2b-f). Postnatal tanycytes closely resemble their adult counterparts^32^, with a cell body lining the ventricular wall, a long basal process, and an endfoot (Fig.2a-b, Extended Data Fig. 1a-d). They also display numerous distinctive protrusions, including spines, irregular swellings, and boutons (Extended Data Fig. 1b-d). However, compared with the adult brain, postnatal tanycytes exhibit a markedly more elaborate morphology, characterized by tortuous processes bearing numerous and elongated spine-like protrusions in their proximal region (Fig. 2b, Extended Data Fig. 1c-d). These complex protrusions are closely associated with neuronal elements, as numerous VGLUT2⁺ presynaptic terminals are detected terminating on them (Fig. 2c, e; Extended Data Fig. 1e)^32,33^. Upon contact with tanycytes, these terminals frequently induced membrane invaginations on the protrusions (Fig. 2e, arrow). In addition to these surface terminations, VGLUT2⁺ terminals are also detected embedded within the tanycytic cytoplasm (Fig. 2d and f, Extended Data Fig. 1e), predominantly in the proximal portion of their processes (Fig. 2d and f). Strikingly, many of these internalized VGLUT2⁺ synaptic elements colocalize with LAMP2⁺ lysosomes (Fig. 2g-h, Extended Data Fig. 1f), suggesting active phagolysosomal degradation. Consistent with this interpretation, these synaptic proteins detected within tanycytes are unlikely to be tanycyte-derived, as publicly available single-cell RNA sequencing datasets show that tanycytes do not express canonical synaptic genes such as *Slc17a7* (Vglut1), *Slc17a6* (Vglut2), *Dlg4* (Psd95), *Homer1*, *Bsn* (Bassoon), *Slc32a1* (Vgat), *Gphn* (Gephyrin), and Gaba-related transcripts during postnatal development (Extended Data Fig. 2). We finally quantified the volume of tanycytic processes occupied by VGLUT2^+^ pre-synaptic elements across postnatal development: the density of VGLUT2⁺ puncta within tanycytic processes peaks at P15 (Fig. 2i), coinciding with the period of synaptic reduction in the mediobasal hypothalamus.

**Figure 2.**
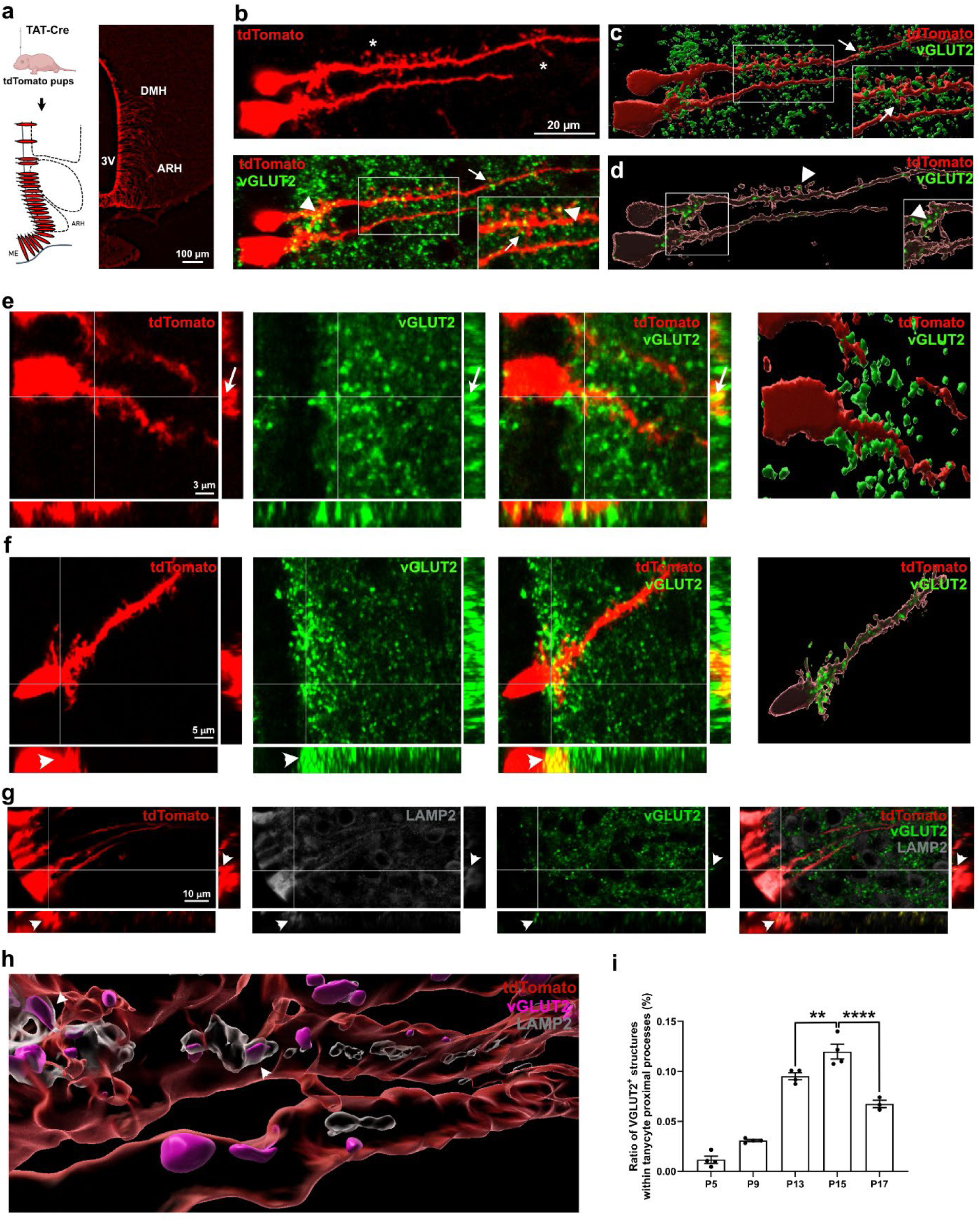
Tanycytes interact and engulf synapses during the postnatal hypothalamic development. **a,** Summary of the analytic workflow: TAT-Cre was infused into the lateral ventricle of tdTomato reporter mice at postnatal day 1 (P1), resulting in tdTomato expression along the third ventricle. In P15 neonates, tdTomato fluorescence (red) filled the entire cytoplasm of tanycytes, allowing visualization of all protrusions along their processes. **b**, High magnification image of tdTomato^+^ tanycytes (red) showing long and thin spines along their process (upper panel, star) and VGLUT2^+^ boutons (green) contacting them (arrows) or engulfed within tanycytic processes (arrowheads). **c**, 3D reconstruction of tanycytes (red) shown in panel b, illustrating VGLUT2⁺ boutons (green) contacting tanycyte processes (arrows). **d**, 3D reconstruction of the interior of tanycytes (gray) shown in panel b, illustrating VGLUT2⁺ boutons (green) engulfed within tanycytic processes (arrowheads). **e**, High magnification image with orthogonal views showing VGLUT2^+^ boutons (green) ending on tanycyte processes (red), forming membrane invagination (arrows), and the corresponding 3D reconstruction of the acquisition (right panel). **f**, High magnification image with orthogonal views showing VGLUT2^+^ boutons (green, arrowheads) engulfed within tanycyte processes (red), and the corresponding 3D reconstruction of the acquisition (right panel). **g**, High magnification image with orthogonal views showing VGLUT2^+^ boutons (green) within LAMP2^+^ lysosomal compartments (gray, arrowheads) within tanycyte processes (red). **h**, 3D reconstruction of the interior of tanycytes (red) showing VGLUT2^+^ boutons (pink) located both outside and within LAMP2^+^ lysosomal compartments (gray). **i**, Quantification of VGLUT2^+^ synaptic elements within tanycyte processes across postnatal development. 3v, third ventricle; ARH, arcuate nucleus; DMH, dorsomedial nucleus; ME, median eminence; Scale bars: as indicated. Data represent mean ± s.e.m.; n = biological replicates (black dots in i); statistical details in Methods. *, p<0.05; **, p<0.01; ***, p<0.001; ****, p<0.0001.

To further characterize tanycyte-synapse interactions and confirm synapse engulfment by tanycytes, we performed sequential array tomography scanning electron microscopy (AT-SEM) and 3D reconstruction of tanycytic processes in the P15 mouse hypothalamus (Fig. 3). These analyses revealed a spectrum of tanycyte-synapse interactions consistent with a stepwise sequence of synapse engulfment and degradation (Fig. 3, Extended Data Fig. 3). In the initial stage, tanycytic processes extend randomly oriented fine spines that closely approach and partially surround synaptic elements (Fig. 3a-b, Extended Data Media 1). During this stage, synapses remain largely intact, with clearly visible presynaptic vesicles and a synaptic cleft (Fig. 3a-b). These interactions occur preferentially with asymmetric (i.e., glutamatergic) synapses, although symmetric (i.e., GABAergic) synapses are also contacted, but less frequently (Extended Data Fig. 3a-e). Tanycytic spines can associate indifferently with presynaptic, postsynaptic, or both pre- and postsynaptic terminals (Extended Data Fig. 3e). Notably, some tanycyte spines can also extend into the synaptic cleft, sometime physically separating the pre- and postsynaptic elements (Fig. 3a-b). Next, the presynaptic element becomes more extensively ensheathed by the tanycytic membrane, forming a nearly complete envelopment while the structural continuity with the axon is still maintained (Fig. 3c-d, Extended Data Media 2). As the interaction progresses, pre-synaptic elements appear entirely internalized within tanycytic processes, with some components, including vesicles and mitochondria, still detectable (Fig. 3e-f, Extended Data Media 2). Finally, synaptic material is observed close to phagosome-like compartments, often lacking distinguishable presynaptic structures, suggesting active degradation (Fig. 3g). The observed time-course and stepwise engulfment are consistent with a phagocytosis-like process (Fig. 3g, Extended Data Fig. 3f) and provide direct evidence of tanycyte-mediated synapse engulfment during this critical postnatal window.

**Figure 3.**
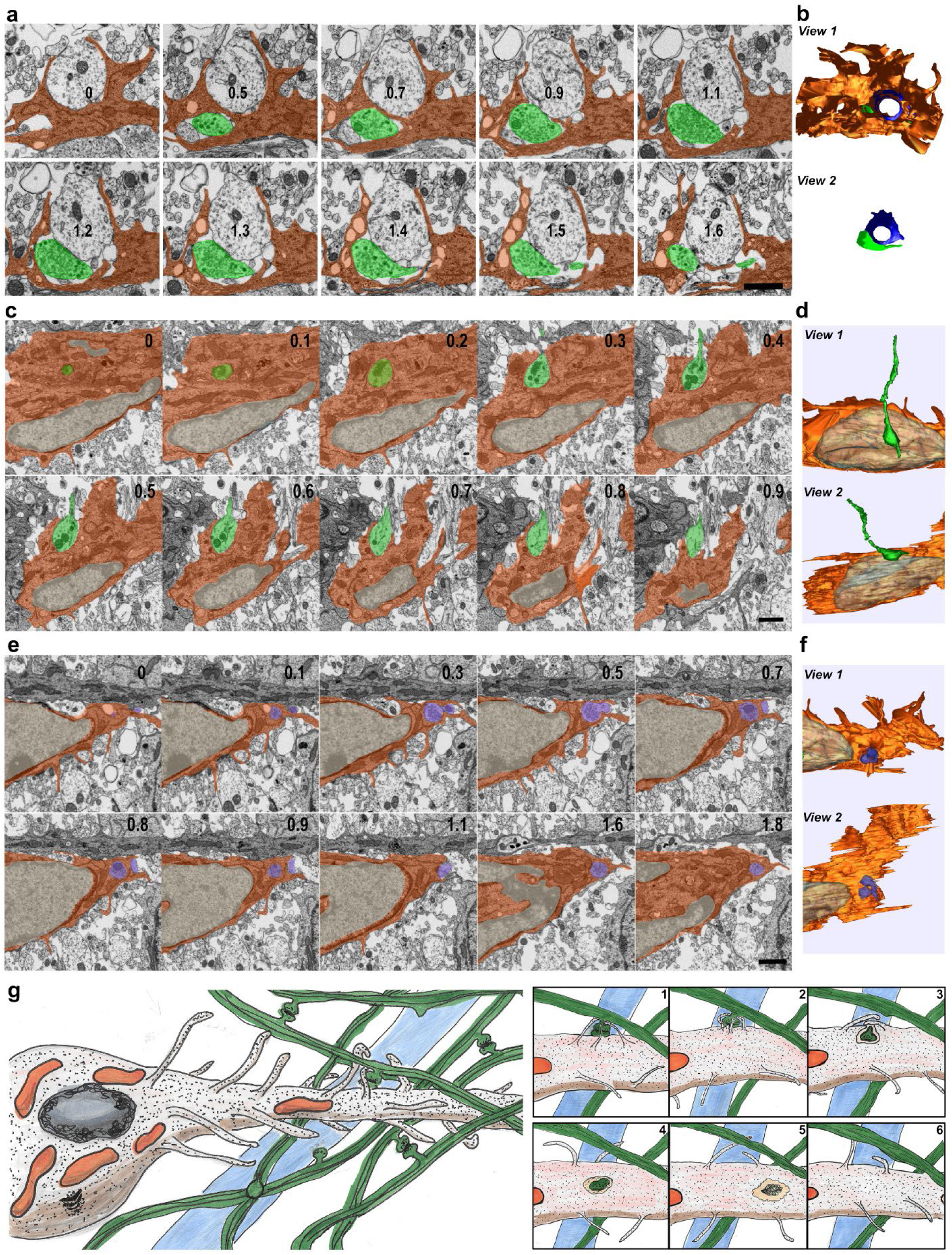
Ultrastructural evidence of synapse engulfment by tanycytes in P15 hypothalamus. **a**, Scanning electron microscopy (SEM) of selected sequential images showing a pre-synaptic element (green) in contact with a tanycytic spine-like extension (orange). Synaptic vesicle and cleft are visible. Numbers inside transversely sectioned tanycyte cell projection indicate the approximate spacing in z direction between the sequential sections in µm. Scale bar 1 µm. **b**, 3D reconstruction obtained by rendering manually traced outlines from the sequential images showed in panel a. Two different views are presented: View 2 corresponds to view 1 without the tanycyte reconstruction. **c**, Scanning electron microscopy (SEM) of selected sequential images showing partial engulfment of a pre-synaptic element (green) by tanycytic processes (orange). Numbers inside transversely sectioned tanycyte cell projection indicate the approximate spacing in z direction between the sequential sections in µm. Scale bar 1 µm. **d**, 3D reconstruction obtained by rendering manually traced outlines from the sequential images showed in panel c. Two different views are displayed, rotated along the x-axis. **e**, Scanning electron microscopy (SEM) of selected sequential images showing full internalization of synaptic material. Numbers inside transversely sectioned tanycyte cell projection indicate the approximate spacing in z direction between the sequential sections in µm. Scale bar 1 µm. **f**, 3D reconstruction obtained by rendering manually traced outlines from the sequential images showed in panel e. Two different views are displayed, rotated along the x-axis. **d**, Schematic illustration summarizing stages of synaptic material interaction with tanycytes. This pseudo-time line represents key steps in the synaptic engulfment, which are supported by our fluorescent and electron microscopy data.

### Tanycytes engulf and eliminate synapses *in vitro*

To further assess tanycytic capacity to engulf and eliminate synapses, we established an *in vitro* phagocytosis assay using primary tanycyte cultures^36^ incubated with fluorescently labeled synaptosomes (*i.e.*, membrane-bound fragments containing pre- and post-synaptic components) isolated from hypothalamic tissue (Fig. 4). Western blot analysis of synaptosome fractions revealed the presence of VGLUT2, PSD-95, and VGAT (Extended Data Fig. 4a-b), as expected^37^. Minimal synaptosome uptake was observed after one hour of incubation (Fig. 4a, c), whereas tanycytes display robust accumulation after four hours (Fig. 4b, c). Tanycyte-engulfed synaptosomes partially colocalize with the lysosome-associated membrane protein 2 (LAMP2) (Fig. 4d), indicating that engulfed synaptic material is trafficked through the phagolysosomal pathway.

**Figure 4.**
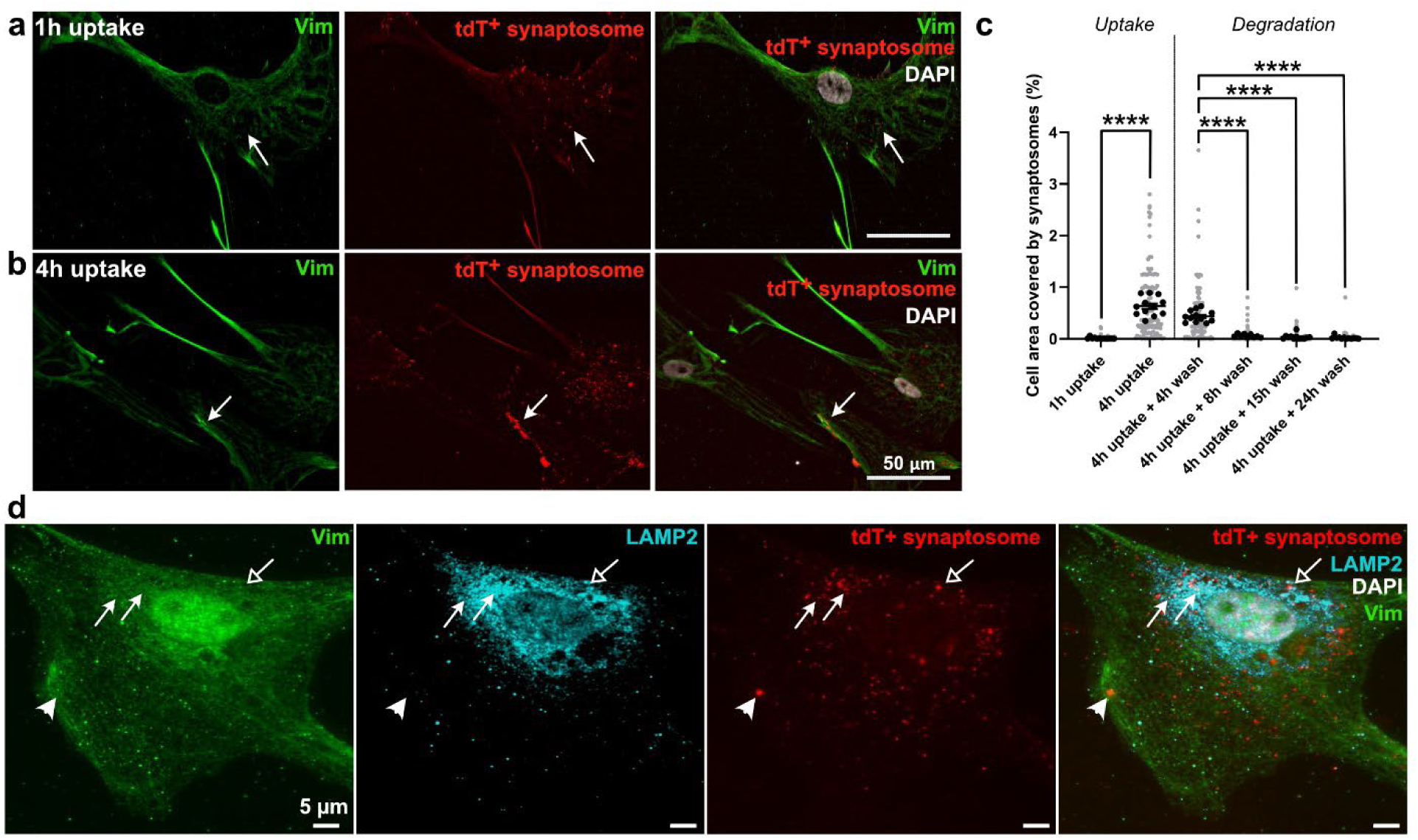
Tanycytes engulf and degrade synaptosomes *in vitro*. **a-b**, Representative images of single tanycytes (green) in primary culture engulfing synaptosome-like structures (red) after 1h (a) and 4h of incubation (b). **c**, Quantification of tanycyte area covered by tdTomato^+^ synaptosomes over time. For uptake, tanycytes were incubated with synaptosomes for 1h or 4h and then fixed. For degradation, tanycytes were incubated with synaptosomes for 4h, washed extensively, and then fixed 4h, 8h, 15h, or 24h later. **d**, High-magnification image of a single tanycyte in primary culture showing synaptosome-like structures (red) located outside (arrowhead), apposed to (empty arrow), or within LAMP2⁺ compartments (arrows) (blue), indicating ongoing degradation. Scale bars: as indicated. Data represent mean ± s.e.m.; n = technical replicates in 4 different cultures (black dots in c), Gray dots in c represent each analyzed tanycytes; statistical details in Methods. *, p<0.05; **, p<0.01; ***, p<0.001; ****, p<0.0001.

To track synaptosome degradation over time, the cultures were washed after four hours of synaptosome exposure and subsequently fixed at 4-, 8-, 15-, and 24-hours post-wash (Fig. 4c). Quantitative analysis revealed a progressive decline in synaptosome-associated fluorescence within tanycytes, with a substantial reduction evident by 8 hours (Fig. 4c). These results confirm that tanycytes not only engulf synaptic material but also actively degrade it via lysosomal processing.

### Tanycytes secrete MFG-E8 as a phagocytic ligand required for synaptosome uptake

To identify candidate molecules involved in tanycyte-mediated synapse engulfment, we analyzed publicly available single-cell RNA-sequencing (scRNA-seq) datasets from the developing and adult hypothalamus^6,35^. Among genes enriched in tanycytes across multiple postnatal stages (P8, P17, P45, and adult brains), Milk fat globule-EGF factor 8 (*Mfge8*) emerged as a strong candidate (Fig. 5a-b, Extended Data Fig. 2). MFG-E8 is a secreted glycoprotein known to facilitate glial-mediated synaptic pruning in other brain regions by bridging phosphatidylserine-exposing targets to integrin receptors on phagocytic cells^16,17^. Genome-wide association studies have further linked the *MFG-E8* gene to BMI-adjusted hip circumference^38,39^, a phenotype notably controlled by the hypothalamus^40^. Among its known receptors, *Itgb3* was not detected in the dataset, whereas *Itgb5* was consistently expressed in tanycytes and microglia across postnatal stages (Fig. 5b, Extended Data Fig. 2), supporting the potential involvement of MFG-E8–ITGB5 signaling axis in synaptic remodeling.

**Figure 5.**
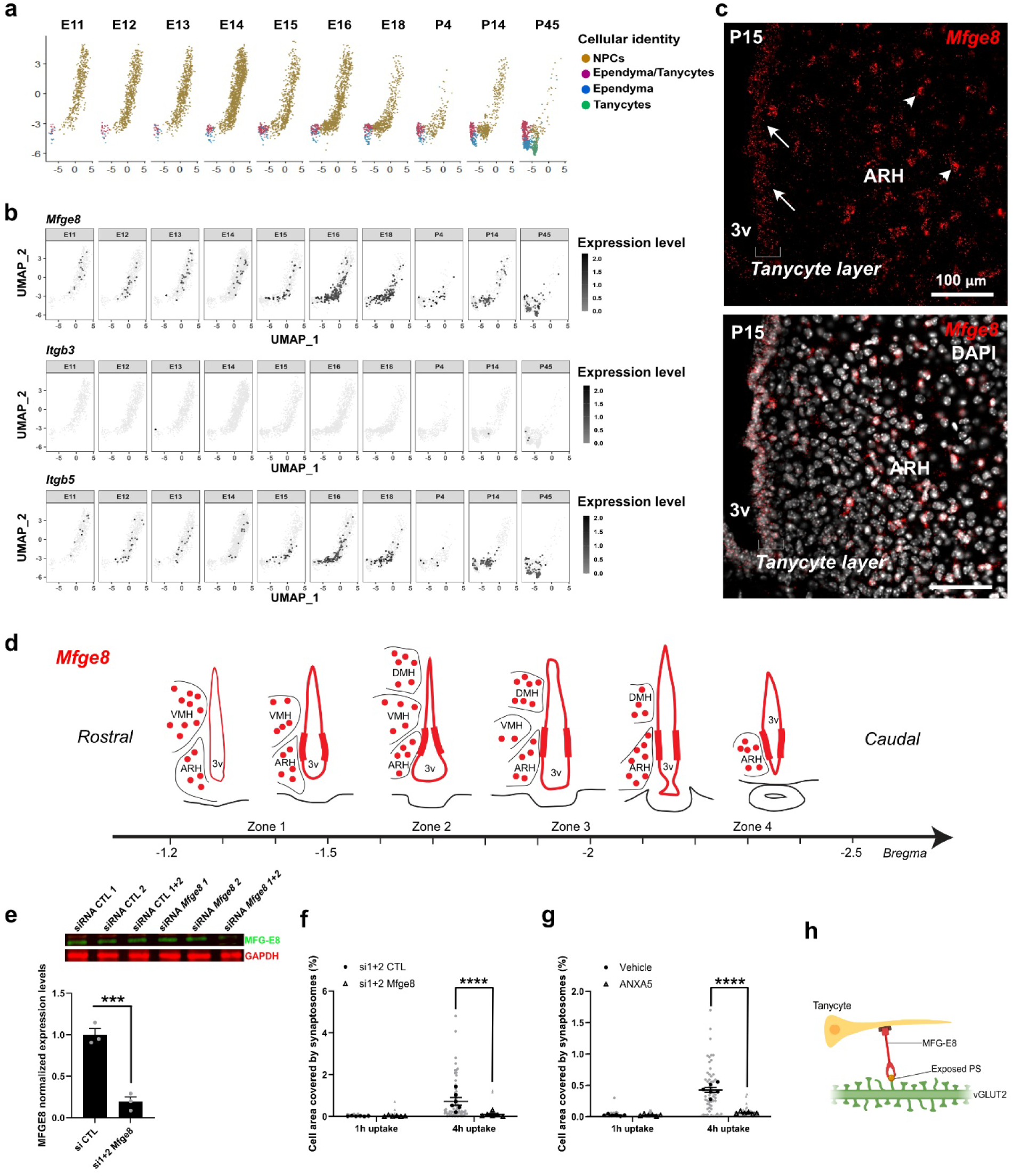
Tanycyte uptake of synapses is *Mfge8*- and phosphatidylserine-dependent. **a,** UMAP displaying the distribution of neural progenitor cells (NPCs, brown), tanycytes (green), and ependymal cells (blue) split by age from E11 to P45. **b,** UMAP showing gene expression (gray to black) of *Mfge8*, *Itgb3*, and *Itgb5* across the developmental period in neural progenitor cells, tanycytes, and ependymal cells, split by age from E11 to P45. Tanycytes and ependymal cells express *Mfge8* and *Itgb5*. **c**, Representative images showing distribution of *Mfge8* mRNAs (red) with DAPI counterstaining (white) along the third ventricle lining the ARH. *Mfge8* is expressed by tanycytes (arrows) and parenchymal cells (arrowheads). **d**, Schematic representation of *Mfge8* expression (red) along the third ventricle on the ventrodorsal and anteroposterior axis based on *in situ* hybridization analysis. **e**, MFGE8 protein expression in primary tanycyte cultures after *Mfge8* siRNA-mediated knockdown. **f,** Quantification of tanycyte area covered by tdTomato^+^ synaptosomes after 1h and 4h of uptake, following synaptosome incubation with or without *Mfge8* knockdown. **g,** Quantification of tanycyte area covered by tdTomato^+^ synaptosomes after 1h and 4h of uptake, following synaptosome incubation with or without ANXA5. **h,** Schematic representation of the MFG-E8 pathway for synaptic pruning by tanycytes. 3v, third ventricle; ARH, arcuate nucleus; ME, median eminence; PS, phosphatidylserine; Scale bars: as indicated. Data represent mean ± s.e.m.; n = technical replicates in 2 different cultures (black dots in c), Gray dots in c represent each analyzed tanycytes; statistical details in Methods. *, p<0.05; **, p<0.01; ***, p<0.001; ****, p<0.0001

To characterize *Mfge8* expression in postnatal tanycytes, RNAscope® in situ hybridization was performed at P15 (Fig. 5c). A strong *Mfge8* signal was observed in the tanycyte layer lining the third ventricle and in parenchymal cells (Fig. 5c), identified as astrocytes based on scRNA-seq data (Extended Data Fig. 2). Along the ventrodorsal axis of the third ventricle, *Mfge8* expression was highest in ARH (α2) tanycytes (Fig. 5d), with no significant variation along the anteroposterior axis (Fig. 5d). In primary tanycyte cultures obtained from P10 rodents, *Mfge8* mRNA was also detected and upregulated fourfold after 24 h of glucose deprivation, indicating responsiveness to metabolic cues (Extended Data Fig. 4c). Proteomic analyses of cell lysates, soluble secretome, and extracellular vesicle fractions from these primary tanycyte cultures^41^ further revealed the presence of MFG-E8 in both the soluble and vesicular fractions, confirming its secretion by tanycytes under basal conditions (Extended Data Fig. 4d).

To assess whether tanycyte-derived MFG-E8 is required for synapse uptake, *Mfge8* expression was silenced in primary tanycyte cultures using siRNAs. Knockdown efficiency was confirmed by a significant reduction in MFG-E8 protein levels when the siRNAs were combined (Fig. 5e). In the synaptosome uptake assay, internalization remained minimal after one hour and was unaffected by *Mfge8* knockdown (Fig. 5f). However, after four hours, tanycytes with reduced MFG-E8 expression exhibited a significant decrease in synaptosome internalization compared with control cells (Fig. 5f), demonstrating that MFG-E8 is required for efficient engulfment of synaptic material by tanycytes.

### Phosphatidylserine recognition enables synaptosome uptake by tanycytes

Given that MFG-E8 facilitates phagocytosis by bridging phosphatidylserine-exposing targets to integrin receptors, we tested whether phosphatidylserine exposure on synaptosomes is necessary for their uptake by tanycytes (Fig. 5g). To do so, primary tanycyte cultures were treated with fluorescent synaptosomes in the presence or absence of Annexin A5 (ANXA5), a protein that binds to and masks externalized phosphatidylserine, thereby preventing its recognition by phagocytic ligands^15^. As in the previous experiment, synaptosome uptake was minimal at 1 hour and not significantly affected by ANXA5 treatment (Fig. 5g). However, after 4 hours, ANXA5 significantly reduced synaptosome engulfment by tanycytes compared to untreated controls (Fig. 5g). These findings confirm that recognition of phosphatidylserine is essential for efficient synaptosome internalization, consistent with the involvement of MFG-E8 as a phosphatidylserine-binding ligand mediating this process (Fig. 5h).

### Tanycyte-specific Mfge8 deletion alters body weight gain and glucose homeostasis

To finally examine the physiological relevance of tanycytic *Mfge8* during postnatal development, we generated tanycyte-specific *Mfge8* knockout (TanMfge8^−/−^) mice by infusing TAT-cre into the lateral ventricle of *Mfge8*^F/F^ mice at P1 (Fig. 6, Extended Data Table 1). *Mfge8*^+/+^ mice receiving the same injection were used as controls. Efficient recombination along the third ventricle was confirmed by a decrease in *Mfge8* mRNA puncta along the ventricular wall facing the ARH (Extended Data Fig. 4e-f).

**Figure 6.**
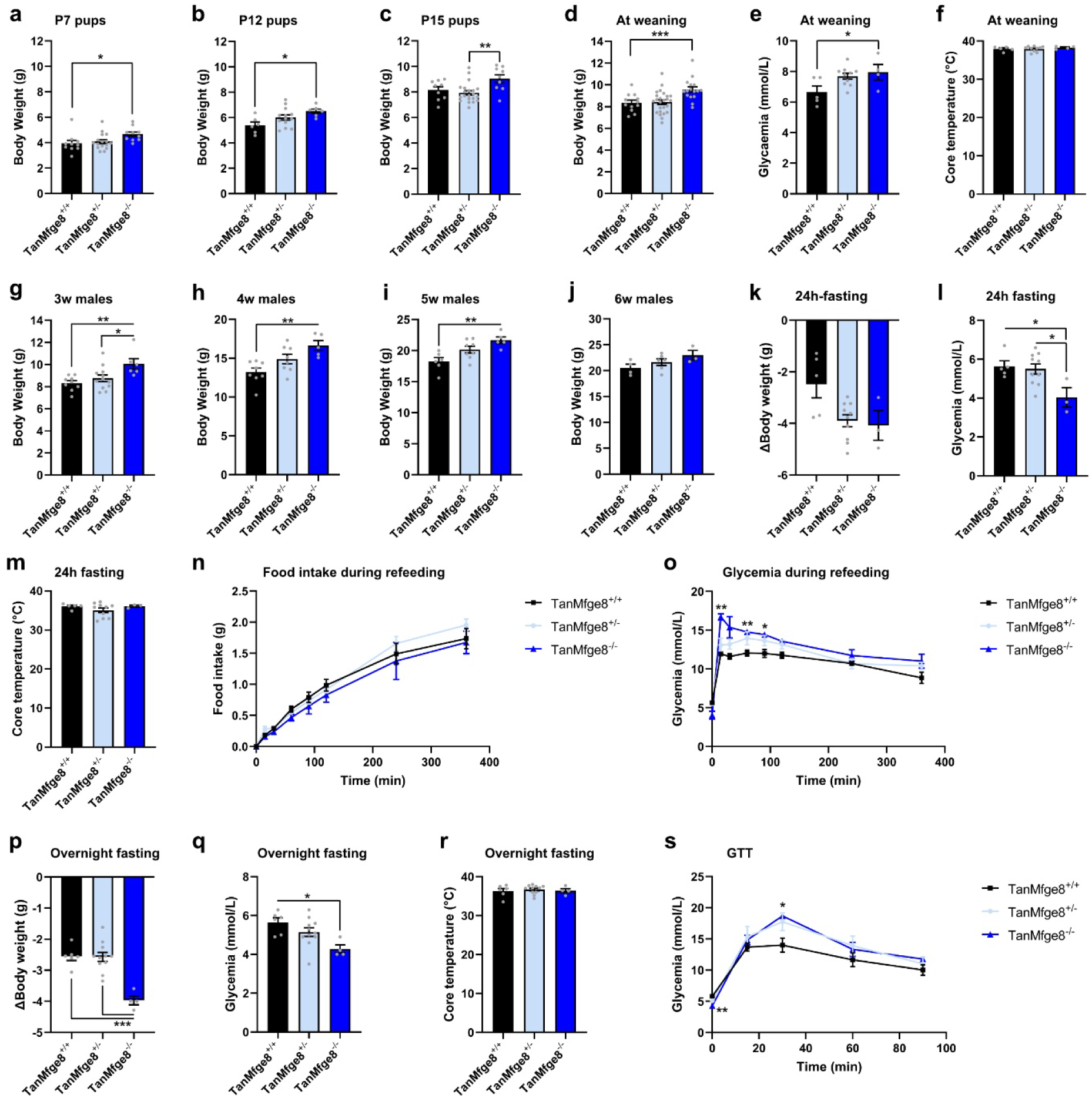
Tanycyte-specific *Mfge8* deletion alters body weight regulation during the postnatal period and impairs glucose homeostasis in adulthood. **a-c,** Body weight in TanMfge8^+/+^, TanMfge8^+/-^, and TanMfge8^-/-^ female and male pups at P7 (a), P12 (b), and P15 (c) following TAT-Cre injection at P1. **d-f,** Body weight (d), glycemia (e), and core body temperature (f) in TanMfge8^+/+^, TanMfge8^+/-^, and TanMfge8^-/-^ male and female pups at weaning following TAT-Cre injection at P1. **g-j,** Body weight in TanMfge8^+/+^, TanMfge8^+/-^, and TanMfge8^-/-^ male mice at 3 (g), 4 (h), 5 (i), and 6 weeks (j) of age following TAT-Cre injection at P1. **k-m,** Delta body weight (k), glycemia (l), and core body temperature (m) in TanMfge8^+/+^, TanMfge8^+/-^, and TanMfge8^-/-^ male mice following 24h-fasting. **n-o,** Cumulative food intake (n) and glycemia (o) in TanMfge8^+/+^, TanMfge8^+/-^, and TanMfge8^-/-^ male mice during refeeding following 24h-fasting. **p-r,** Delta body weight (p), glycemia (q), and core body temperature (r) in TanMfge8^+/+^, TanMfge8^+/-^, and TanMfge8^-/-^ male mice following overnight fasting. **s,** Glycemia in TanMfge8^+/+^, TanMfge8^+/-^, and TanMfge8^-/-^ male mice during glucose tolerance test following overnight fasting. Data represent mean ± s.e.m.; n = biological replicates (gray dots); statistical details in Methods. *, p<0.05; **, p<0.01; ***, p<0.001; ****, p<0.0001

Body weight (BW) monitoring revealed a significant increase in male and female TanMfge8^−/−^ mice compared to controls, detectable from P7 and persisting until weaning (Fig. 6a-f). At weaning, male TanMfge8^−/−^ mice display higher BW, which remains elevated up to 5 weeks of age (Fig. 6g-j, Extended Data Table 1). This is accompanied by higher glycemia but no change in body temperature (Fig. 6e-f). After 6 weeks, the BW of control mice reaches that of the knockout mice.

In adulthood, TanMfge8^−/−^ mice exhibit alterations in body weight regulation and glycemic control. Notably, 24h-fasting tends to induce a more pronounced BW loss and leads to deeper hypoglycemia (Fig. 6k-l), without affecting core body temperature (Fig. 6m). During the refeeding phase, food intake amount and pattern are similar between groups (Fig. 6n). However, blood glucose levels show a sharper rise at the onset of refeeding in TanMfge8^−/−^ mice (Fig. 6o), suggesting a dysregulation in glucose handling. Similarly, overnight fasting results in greater weight loss and more pronounced hypoglycemia in TanMfge8^−/−^ mice (Fig. 6p-q), again without changes in core body temperature (Fig. 6r). During the glucose tolerance test (GTT), TanMfge8^−/−^ mice display a more pronounced hyperglycemic response (Fig. 6s), indicating potential impairments in glucose clearance.

## DISCUSSION

Here, we uncover a previously unrecognized role for tanycytes as active participants in synaptic remodeling during postnatal development. This process is mediated by the secreted ligand MFG-E8, which bridges phosphatidylserine-exposing presynaptic terminals to promote their engulfment. Tanycyte-specific deletion of *Mfge8* during early postnatal life leads to increased body weight at weaning and impaired glycemic control in adulthood, indicating that tanycyte-mediated synapse elimination is essential for the proper maturation of hypothalamic circuits governing metabolic regulation. Together, these findings expand the functional repertoire of tanycytes beyond their established roles in neurogenesis^35,42–45^, transport^28,29,46^, and signaling^31,41^, and position them as key sculptors of neuronal connectivity in the developing hypothalamus.

The establishment of hypothalamic circuits governing energy balance relies on tightly coordinated neurodevelopmental processes, including neurogenesis, axonal projections, synaptogenesis, and synaptic refinement^3,8^, that extend into the third postnatal week^47^. During this critical window, peripheral hormonal cues such as leptin^48^, insulin^49^, and ghrelin^50^, dynamically interact with intrinsic developmental programs to tune neuronal connectivity within emerging hypothalamic circuits^51–53^. Although microglia are well-established mediators of synaptic pruning^54^ in other brain regions^12,13,55^, their contribution in hypothalamic circuit formation has only recently been recognized^18,56,57^ and appears heterogeneous and cell-type specific^58^, with distinct effects on AgRP^18,56,57^ *versus* POMC neurons^56^. Our findings now add tanycytes to this picture, identifying them as an additional and previously unrecognized glial population engaged in synaptic remodeling. This discovery challenges the canonical view that microglia are the sole executors of developmental synapse elimination in the hypothalamus and introduces a new paradigm in which ependymal cells contribute to shaping neural circuits underlying metabolic homeostasis.

While lysosomal degradation by tanycytes has been reported in the context of lipid processing in the mature brain^59^, no prior evidence has indicated phagocytic activity during development. However, this newly identified function parallels that of their evolutionary ancestors, radial glia, which actively clear cellular debris, apoptotic cells, and synaptic material in other brain regions and species through specialized and context-dependent mechanisms^60,61^. Tanycyte phagocytic activity itself appears to be developmentally regulated, peaking around postnatal day 15, a developmental window that coincides with tanycyte maturation^6,34^ and major hormonal^51^ and behavioral transitions^3^ in rodents. This temporal specificity suggests that tanycyte-mediated pruning is both developmentally programmed and metabolically tuned, underscoring the need to further explore the cellular and molecular diversity of synaptic pruning mechanisms in the hypothalamus in a context-dependent manner.

Mechanistically, our data identify MFG-E8 as a critical molecular mediator of tanycyte phagocytosis. MFG-E8 functions as an opsonin that bridges “eat-me” signals such as phosphatidylserine on neuronal membranes to integrin receptors on glial cells^62^. Well characterized in astrocyte–microglia communication in cortical^16^ and hippocampal^63^ synapse elimination, MFG-E8 levels have been associated with increased synapse loss in neurodegenerative diseases whereas both astrocytic *Mfge8* knockdown^63^ and pharmacological blockade of MFG-E8^16^ prevent microglial engulfment and mitigate synaptic degradation. Here, we demonstrate that tanycytes use this same molecular toolkit for autonomous synaptic engulfment, reinforcing the concept that MFG-E8 serves as a general mediator of glial-driven synapse elimination across diverse cell types and brain regions. Importantly, we also find that MFG-E8 expression in tanycytes is sensitive to glucose availability, implicating a coupling between nutrient status and structural remodeling. While this appears to contrast with previous reports showing that serum MFG-E8 decreases during fasting and rises upon refeeding^64,65^, with insulin and high glucose promoting MFG-E8 redistribution to the cell surface in muscles^64^ and secretion via extracellular vesicles by endothelial cells^66^, such discrepancies reflect cell type-specific and context-dependent regulatory mechanisms. Together, these findings suggest that MFG-E8 signaling is finely tuned by the metabolic state and may operate differently across tissues. In the hypothalamus, such metabolic gating of tanycyte phagocytic activity positioning these cells as integrators of organism’s metabolic state, ensuring that hypothalamic circuit refinement proceeds in synchrony with metabolic demands.

Tanycytes secrete MFG-E8 both in the soluble secretome and via extracellular vesicles. While this supports autocrine synaptic pruning, the expression of ITGB5, the receptor for MFG-E8, by neighboring microglia and astrocytes^67^ raises the possibility of paracrine signaling, potentially coordinating synapse engulfment across multiple hypothalamic glial populations. Indeed, in other brain regions, MFG-E8 mediates astrocyte–microglia crosstalk during synapse elimination^63^. Furthermore, tanycyte–microglia communication has been observed in the adult brain under neuroinflammatory conditions^41^, suggesting that similar interactions could contribute to developmental synaptic pruning. Developmental parallels with radial glia, which cooperate with microglia to clear cellular debris, further support this possibility^68,69^. Elucidating whether tanycytes, astrocytes, and microglia act in parallel, redundantly, or cooperatively will be essential for fully mapping the cellular networks governing hypothalamic circuit maturation.

Finally, our physiological data indicate that tanycyte-mediated synaptic pruning plays a critical role in the regulation of energy balance. Mice lacking *Mfge8* specifically in tanycytes during early postnatal life exhibit excessive weight gain at weaning and impaired glycemic control in adulthood, revealing a lasting impact of early-life tanycyte activity on energy homeostasis. These findings are consistent with studies in peripheral tissues, where MFG-E8 influences glucose homeostasis by modulating insulin signaling^65^. Notably, elevated serum MFG-E8 levels have been reported in obese and diabetic mice, where it promotes fat absorption and insulin resistance via integrin-dependent signaling^70,71^.^70,71^ Our data now extend this concept to the central nervous system, suggesting that MFG-E8 may exert dual roles, both systemic and central, in coordinating metabolic balance. Whether these central effects are mediated solely through structural synaptic remodeling or also involve additional signaling functions of MFG-E8 remains an important question for future investigation.

Collectively, this study identifies a novel role for tanycytes in the elimination of presynaptic terminals during postnatal hypothalamic development via the MFG-E8 pathway, redefining them as active participants in the structural maturation of hypothalamic circuits. More broadly, these findings expand the concept of glial diversity, revealing that developmental synaptic pruning is region-specific and adapted to local circuit demands. Understanding how glia interactions evolve across development will be critical to deciphering how hypothalamic circuits governing energy balance are established—and how their disruption may contribute to metabolic disease.

## Supporting information

Extended Data Media 1

Extended Data Media 2

## RESOURCE AVAILABILITY

### Lead contact

Requests for further information and resources should be directed to and will be fulfilled by the lead contact.

### Materials availability

This study did not generate new unique reagents.

### Data and code availability

This study did not generate new codes or datasets.

## ACKNOWLEDGEMENTS

This work was supported by the Swiss National Science Foundation (320030-231343). F.L. is supported by the European Research Council Starting Grant (TANGO, No. 948196), the Novartis Foundation for medical-biological research, and the Department of Biomedical Sciences at the University of Lausanne. The authors also thank the Animal Facility (UNIL), the Protein Analysis Facility (PAF-UNIL), the Cellular Imaging Facility (CIF-UNIL), and the Electron Microscopy Facility (EMF-UNIL).

## AUTHOR CONTRIBUTION

R.D. designed and performed experiments, analyzed and interpreted the data, produced the figures, and wrote the manuscript. I.K., C.K.G., and T.D. performed experiments and analyzed the data. D.L.-R. performed bioinformatic analysis. F.L. designed experiments, oversaw research, analyzed and interpreted the data, produced the figures, and wrote the manuscript.

## SUPPLEMENTAL INFORMATION TITLES AND LEGENDS

**Extended Data Figure 1.**
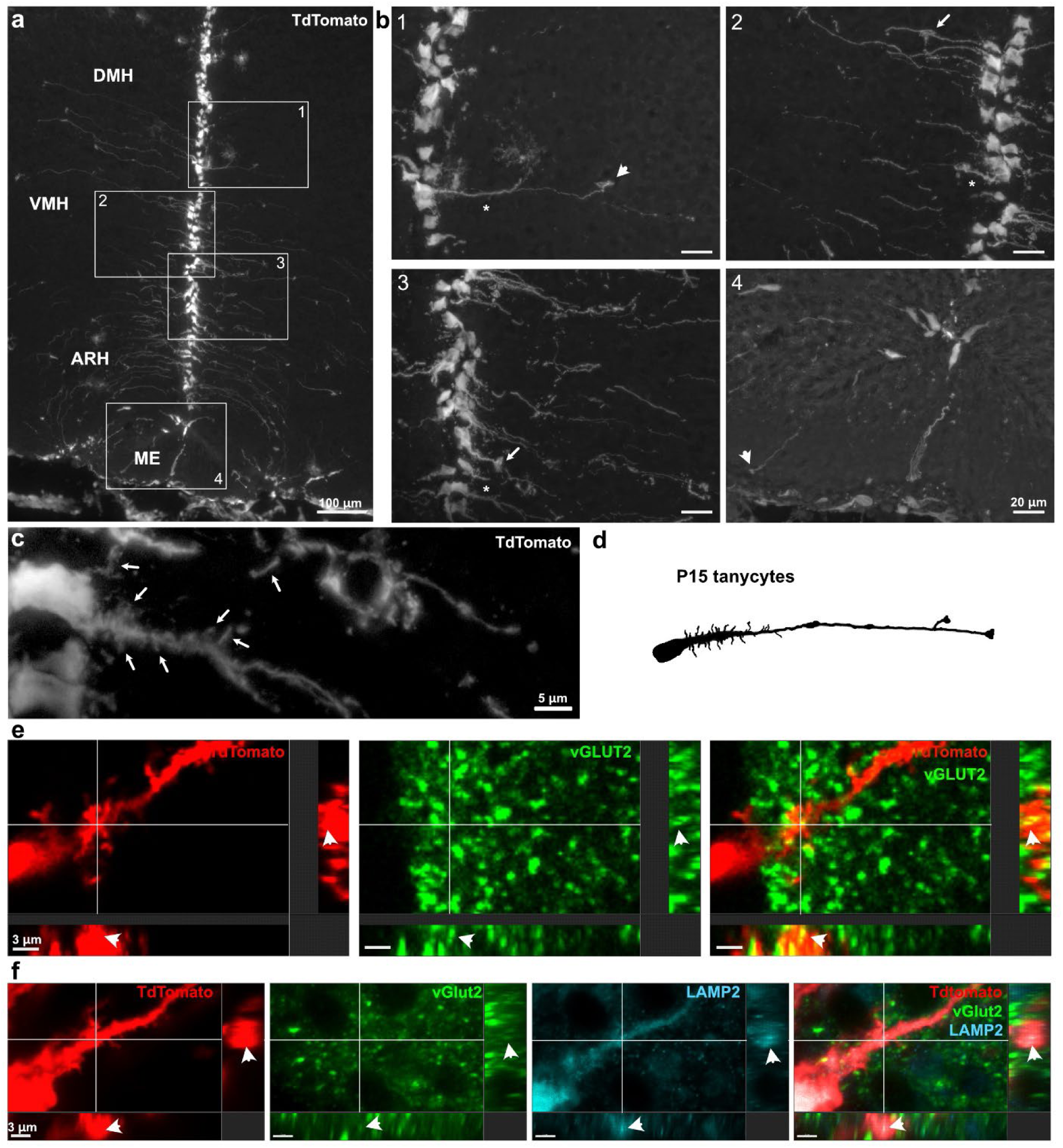
Postnatal tanycyte process morphology observed along the third ventricle in p15 mice. **a**, Representative image (×10) showing the distribution of tdTomato (gray) at P15 along the third ventricle in coronal section following TAT-Cre injection at P1. **b**, Representative z-stack images (×63) showing inserts (from 1 to 4) present in panel a. Protrusions are observed in tanycyte processes lining the lateral wall of the 3V, including spines (stars), swelling (arrows), and boutons (arrowheads). Pictures (d–k) are the maximal intensity projections of the z-stack acquisition. **b**, Numeric zoom of insert 3 to visualize tanycyte spines (arrows). **c,** Schematic representation of P15 tanycytes facing hypothalamic parenchyma (*i.e.*, ARH, VMH, DMH). **d**, High magnification image with orthogonal views showing VGLUT2^+^ boutons (green, arrowheads) engulfed within tanycyte processes (red). **e**, High magnification image with orthogonal views showing VGLUT2^+^ boutons (green, arrowheads) within LAMP2^+^ lysosomal compartments (blue) within tanycyte processes (red). 3V, third ventricle; ARH, arcuate nucleus; DMH, dorsomedial nucleus; ME, median eminence; VMH, ventromedial nucleus. Scale bars, as indicated.

**Extended Data Figure 2.**
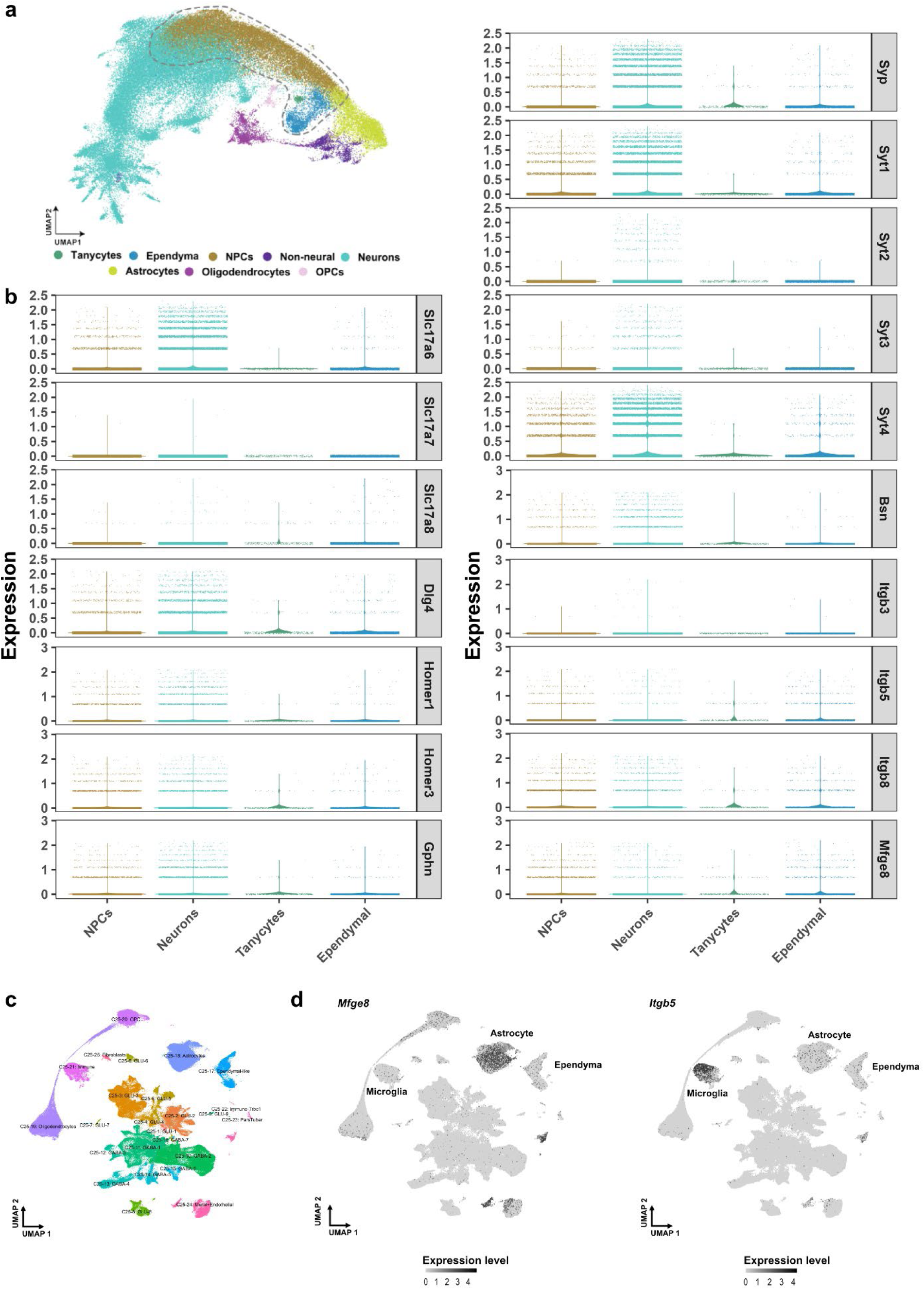
Transcriptomic profiling of hypothalamic neural cell population in the postnatal mouse brain. **a**, UMAP projection showing the integrated developmental scRNAseq datasets from hypothalamic and diencephalic explants between E11 and P45 (excluding E10 and P8 datasets) from Kim *et al.*^6^ (clustering used in Lopez-Rodriguez *et al.*^34^). Colors represent different cell types: tanycytes (green), ependymal cells (blue), neural progenitor cells (NPCs; brown), non-neural cells (purple), neurons (cyan), astrocytes (yellow-green), oligodendrocytes (pink), and oligodendrocyte precursor cells (OPCs; light pink). **b**, Violin plots showing gene expression profiles in neural progenitor cells (NPCs), neurons, tanycytes, and ependymal cells. Panels highlight genes associated with synaptic vesicular trafficking and structure markers (e.g., *Slc17a6, Slc17a7, Slc17a8, Dlg4, Homer1, Homer3, Gphn, Syp, Syt1, Syt2, Syt3, Syt4, Bsn*), cell adhesion and signaling (e.g., *Itgb3, Itgb5, Itgb8*), and our gene of interest (e.g., *Mfge8*). **c**, UMAP projection showing the adult mouse Hypomap integrated scRNAseq datasets^67^, colored by cell types. **d**, UMAP projection showing the gene expression of *Mfge8* and *Itgb5* across the dataset. Gray scale corresponds to the expression level.

**Extended Data Figure 3.**
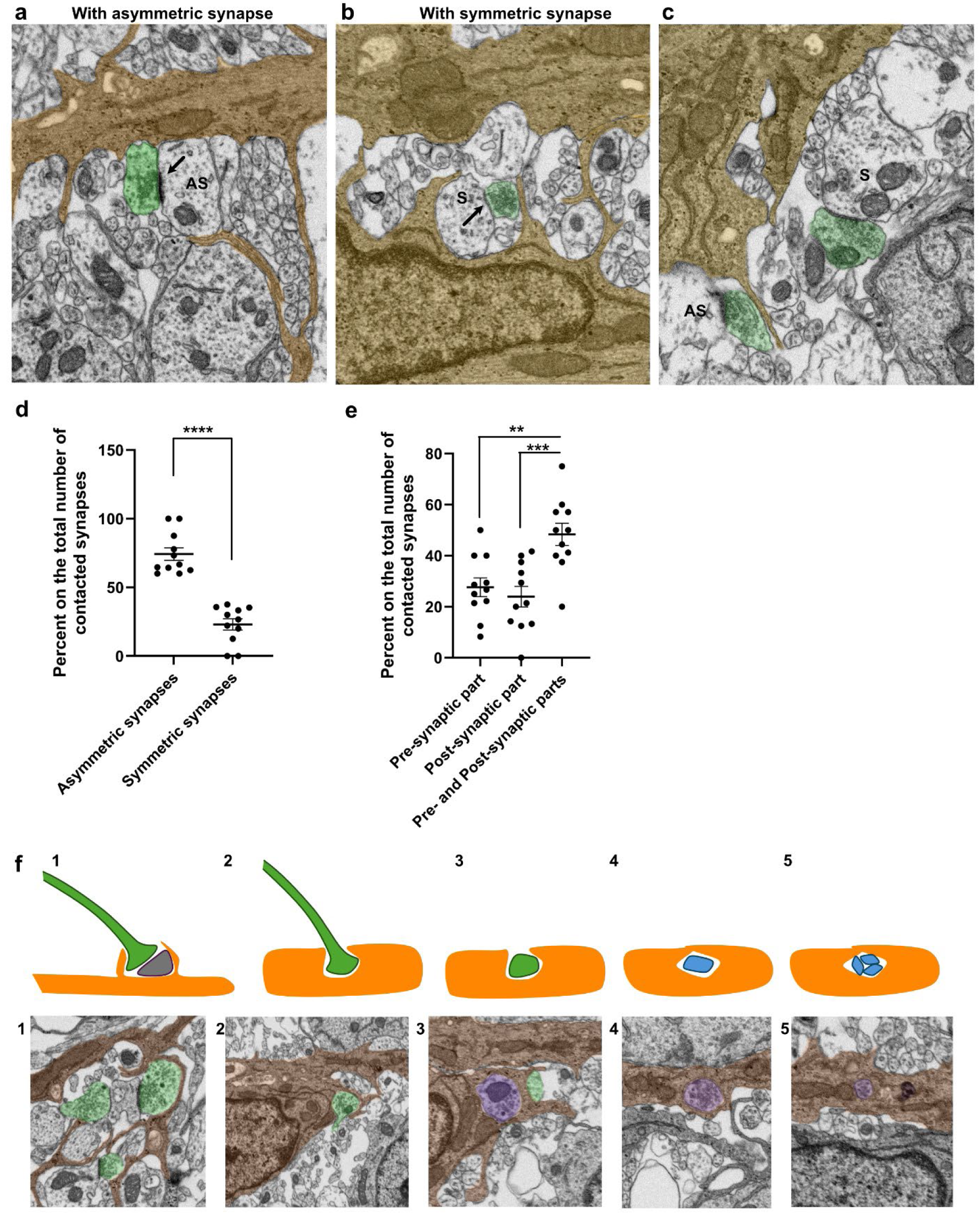
Tanycytes contact both symmetric and asymmetric synapses. **a**, Scanning electron microscopy (SEM) images showing an asymmetric synapse in contact with a tanycytic spine-like extension (orange). The presynaptic part is colored green. The synaptic cleft is large and asymmetrical (arrow). **b**, Scanning electron microscopy (SEM) images showing a symmetric synapse in contact with a tanycytic spine-like extension (orange). The presynaptic part is colored green. The synaptic cleft is thin and symmetrical (arrow). **c**, Scanning electron microscopy (SEM) images showing an asymmetric synapse (AS) in contact with a tanycytic spine-like extension (orange), while a symmetric synapse (S) remains untouched by tanycytes. The presynaptic part is colored green. **d-e,** Percentage of tanycyte interactions with symmetric *vs.* asymmetric synapses (d) and with pre-synaptic, post-synaptic, *vs.* both pre- and post-synaptic parts (e). A total of 11 tanycytes were analyzed, with an average of 9.73 ± 1.35 synapses interacting with each tanycyte. **f**, Schematic representation with corresponding scanning electron microscopy (SEM) images illustrating the successive stages of synaptic material interaction with tanycytes.

**Extended Data Figure 4.**
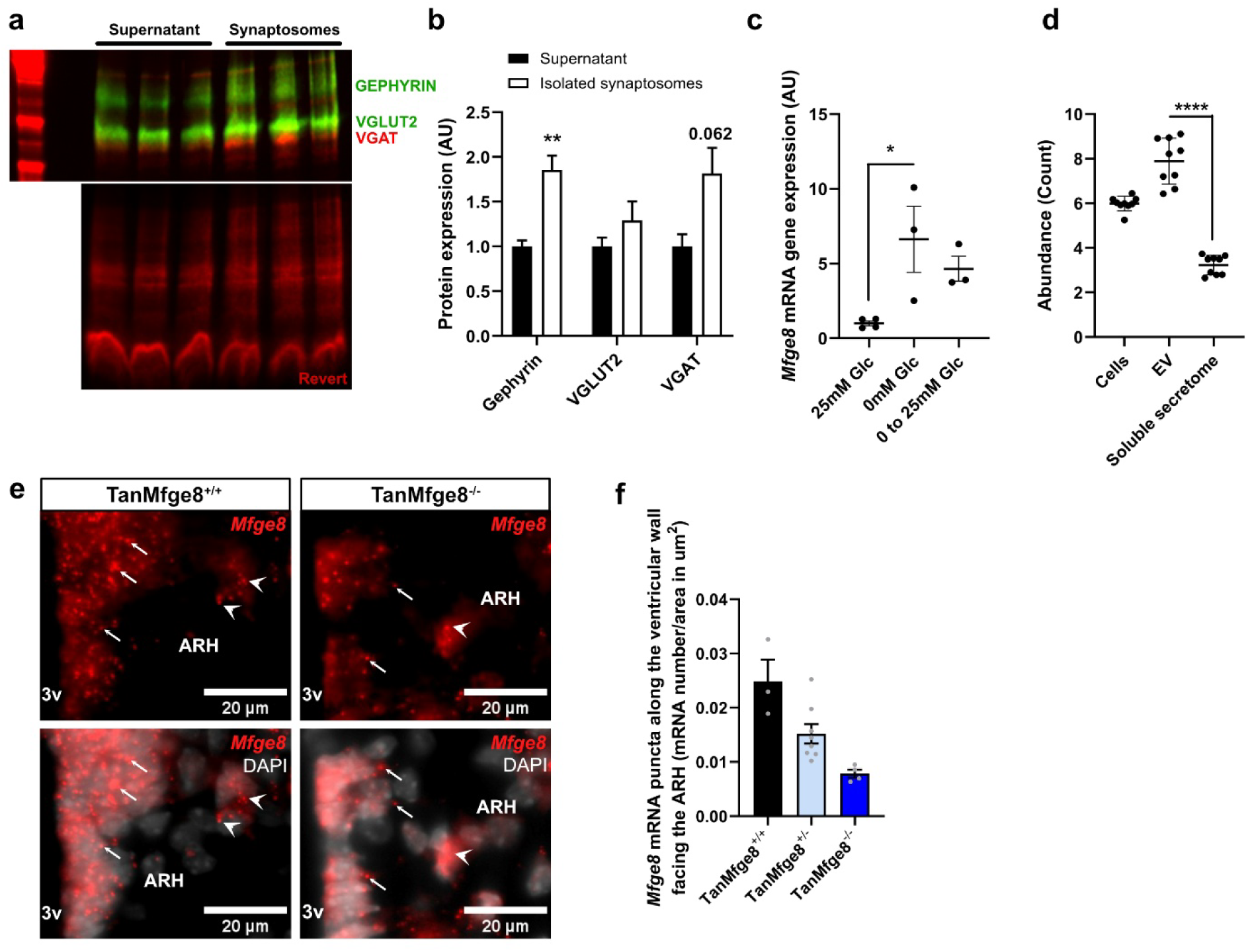
*Mfge8* expression in tanycytes. **a-b,** Western blot analysis of hypothalamic synaptosome preparations demonstrating the presence of VGAT (red), VGLUT2 (green), and Gephyrin (green) (a). The synaptosome isolation protocol induces an enrichment of synaptic material in the final product (b). **c,** *Mfge8* expression measured by qPCR in primary tanycyte cultures following glucose treatment after 24 h of glucose deprivation (n = 3/4 per condition from one culture). **d,** MFG-E8 levels in the cellular, extracellular vesicle, and soluble fractions of tanycyte-conditioned medium, as determined by proteomic analysis. **d-e,** Representative images (e) and quantification (f) of *Mfge8* mRNA along the ventricular wall facing the ARH in CTL *versus* TanMfge8^−/−^ mice at P15.

**Extended Data Media 1. Overview of tanycytic-synaptic interactions: tanycyte spines.**

Whole-mount 3D reconstruction showing tanycytes and surrounding neuronal synapses in the ARH, corresponding to Figure 3 a-b.

**Extended Data Media 2. Overview of tanycytic-synaptic interactions: engulfed synapses.**

Whole-mount 3D reconstruction showing tanycytes and surrounding neuronal synapses in the ARH, corresponding to Figure 3 c-f.

**Extended Data Table 1. Mfge8^+/+^ and ^-/-^ mouse physiology after weaning.**

**Extended Data Table 2. Methods.**

## METHODS

### Mice and genotyping of transgenic animals

C57Bl/6J mice, Rosa26-floxed stop tdTomato mice (initially obtained from Charles River), CamKIIa^cre/+^;Rosa-fl-STOP-TdTomato mice^48^, Mfge8-floxed mice (GemPharmatech), and Sprague-Dawley rats were used in this study. Male and female mice were put together around 5 pm, and the presence of a vaginal plug was checked on the following day (before 9 am). The day of birth was regarded as postnatal day 0 (P0). Litters were standardized to 5–6 pups per dam. Biopsies were collected for genotyping, and DNA extraction was performed using the HotShot method. PCR amplification was performed using KAPA2G Fast ReadyMix (KAPA Biosystems, KK5103) following the manufacturer’s instructions. The primers (5ʹ-3ʹ) used in this study are available on request. All animal procedures were performed at the University of Lausanne and were reviewed and approved by the Veterinary Office of the Canton de Vaud (VD3825, VD3731, and VD3731×1).

### Intracranial injections on P1 pups

To recombine DNA in tanycytes, TAT-CRE fusion protein (MERCK, SCR508; 2 μl; diluted in saline solution at 1:3 or 1:2) was infused into the lateral ventricle of P1 pups. Injection sites were visualized through the translucent skin using anatomical landmarks, including bregma, lambda, and the eyes. Injections were placed either at approximately two-fifths of the distance between lambda and each eye, or 1mm lateral to the sagittal suture midway between bregma and lambda: both approaches reliably target the lateral ventricle^72^.

### Mouse physiology

Body weight was measured every two to three days from birth until weaning using a precision scale. After weaning, body weight, blood glucose, and core body temperature were monitored weekly between 8 and 9 a.m. Blood glucose levels were assessed via tail vein sampling using a Contour NEXT glucose monitor. Body temperature was recorded using a rectal thermometer (BIOSEB, BIO-TK8851). For the fasting-refeeding paradigm, mice were fasted for 24 hours, then isolated for 1 hour before the experiment (starting at 9 a.m.). Food intake, glycemia, and body temperature were measured at different time points during the refeeding period.

For the glucose tolerance test, mice were fasted overnight. Body weight and baseline glycemia were measured at 5 p.m. on the day before the test and again at 9 a.m. before glucose administration. Mice then received an intraperitoneal injection of D-glucose (2g/kg of body weight). Blood glucose was measured from the tail vein at different time points.

### Tissue collection

For immunohistochemistry and *in situ* hybridization on fixed tissue, mice were anesthetized with pentobarbital and perfused transcardially with 0.9% saline, followed by an ice-cold solution of 4% paraformaldehyde in 0.1 M phosphate buffer (pH 7.4). Brains were quickly removed, postfixed in the same fixative for 2h at 4°C, and immersed in 20% sucrose in 0.1M phosphate-buffered saline (PBS) at 4°C overnight. Brains were finally embedded in ice-cold OCT medium (optimal cutting temperature embedding medium, Tissue Tek, Sakura) and frozen on dry ice or in liquid nitrogen-cooled isopentane.

For electron microscopy, P15 mice were anesthetized with pentobarbital and perfused transcardially with 0.9% saline, followed by an ice-cold solution of 4% paraformaldehyde/5% glutaraldehyde in 0.1 M phosphate buffer (pH 7.4). Brains were quickly removed and fixed in an ice-cold solution of 4% paraformaldehyde/5% glutaraldehyde in 0.1 M phosphate buffer (pH 7.4) for 24h. Two hundred µm-thick hypothalamic slices were cut using vibratome and postfixed in the same fixative for 24h. Afterward, the samples were incubated in 2% (wt/vol) osmium tetroxide and 1.5% (wt/vol) K4[Fe(C.N.)6] in 0.1 M phosphate buffer on ice for one hour, and dehydrated at the end of standard gradual dehydration cycles in ethanol. Samples were flat embedded in an Epon-Araldite mix.

### Immunohistochemistry

Brains were cut using a cryostat into 20-μm-thick coronal sections and processed for immunohistochemistry as described previously^73^. Briefly, the slide-mounted sections were 1) incubated in a boiling 10 mM Citrate Buffer solution, pH 6.0, for 12 minutes; 2) blocked for 1 hour using a solution containing 2% normal goat serum and 0.3% Triton X-100; 3) incubated overnight at 4°C with primary antibodies followed by two hours at room temperature with a cocktail of secondary Alexa Fluor-conjugated antibodies (1:500, Molecular Probes, Invitrogen); 4) counterstained with DAPI (1:10,000, Molecular Probes, Invitrogen), and 5) coverslipped using Mowiol (Calbiochem, La Jolla, CA). All primary and secondary antibodies used are listed in Extended Data Table 2.

### In situ hybridization

Fixed frozen brains were cut using a cryostat into 20-μm-thick coronal sections and processed for RNAscope® in situ hybridization following the manufacturer’s instructions (ACD). Slide-mounted sections were first 1) incubated in a boiling 1X Target retrieval solution for 5 minutes, and 2) incubated at 40°C with Protease III solution for 30 minutes. The sequential hybridizations were then performed following the manufacturer’s instructions. The probes used in this study are reported in Extended Data Table 2.

### Fluorescent microscopy

Images were acquired with either a Leica Thunder Imaging System, a Nikon i90 microscope, or a confocal (Zeiss LSM 780, 800, and 900). Frames were collected stepwise over a defined z-focus range corresponding to all visible fluorescence within the section: multiple-plane frames were collected at a step of 1 µm while using x20, x40, or x63 objective (between 4 and 35 frames per image). All images were then saved in .lif or .nds, processed to get orthogonal and maximal intensity projections, and finally exported in .tiff for the processing steps (i.e., adjust brightness and contrast, change colors, and merge images using ImageJ).

### Electron microscopy

Flat-embedded samples were mapped using binoculars to map the precise region of interest at the basal part of the ventricle. Polymerized flat blocks were trimmed using a 90° diamond trim tool and sequential sections of 100 nm were generated using a 35° ATC diamond knife (Diatome, Biel, Switzerland) mounted on a Leica UC6 microtome (Leica, Vienna). These arrays of sections covered the span of the ROI and were transferred on pieces of silicon wafer^74^. Wafers were analyzed using FEI Helios Nanolab 650 scanning electron microscope (Thermo Fischer, Eindhoven). Wafers were laterally screened to target the relevant sections, and the ventricle area was using the following settings: M.D. detector, accelerating voltage 2kV, current 0.8nA, dwell time 4-6µs. After the initial ROI localization, relevant areas were re-imaged using higher resolution parameters, collected manually using MAPs program (Thermo Fischer, Eindhoven) (ref). For EM acquired serial sections images were aligned into a stack, using IMOD program (REF). 3dmod program was used for manual rendering of the datasets. For each section, manual tracking of the outlines was performed and followed by 3D reconstruction to visualize the special interactions between different cells. Pseudo-coloring was maintained to mimic the fluorescent convention.

### Primary tanycyte cultures

Hypothalamic MEs were isolated from 10-day-old (P10) pups (Wistar rat, Janvier Laboratory) for tanycytes primary culture, as previously described^36^. Briefly, after decapitation and collection of the brain, MEs were dissected and crushed on 40-uM nylon mesh (Sefar America). Dissociated cells were cultured in DMEM high glucose (Gibco, cat no 419666052), 10% pen-strep, 2% L-glutamine, supplemented with 10% (v/v) donor calf serum at 37°C under a humid atmosphere of 5% CO2. The culture medium was changed after 3 days of culture and subsequently every 2 days. On reaching confluence, tanycytes were seeded in 24-well plates on poly-L-lysine-coated glass coverslips for immunocytochemistry, uncoated plastic 12-well plates for gene expression, and uncoated plastic 6-well plates for protein expression analysis.

### Electroporation

Proliferating tanycytes were electroporated with siRNAs against Mfge8 or a control siRNA (30pmol) (see Table 1), using 4D-Nucleofector X Unit EA (LZ-AAF-1002X, Lonza) and the P3 primary cell 4D-Nucleofector X solution (LZ-V4XP-3024, Lonza). Electroporated cells were plated in DMEM high glucose (Gibco, cat no 419666052), 10% pen-strep, 2% L-glutamine, supplemented with 10% (v/v) donor calf serum. 24 hours later, cells were trypsinized and plated on poly-L-lysine-coated glass coverslips for immunocytochemistry or uncoated plastic 12-well plates for gene expression, and uncoated plastic 6-well plates for protein expression analysis.

### Isolation of synaptosomes and synaptosome engulfment assay

Synaptosomes were isolated from adult CamKIIa^cre/+^; Rosa-fl-STOP-TdTomato^75^ hypothalamic tissue using Syn-PER® Synaptic Protein Extraction Reagent (Cat#87793, Invitrogen) according to the manufacturer’s instructions. The synaptosomes-containing pellet was resuspended in 5% DMSO in Syn-PER® to obtain a concentration of 6.5 μg/ml. Aliquots were stored at -80 °C. Primary tanycytes were incubated for 1h or 4h with synaptosomes (65 μg/ml in DMEM F12, no serum) and either fixed, stained, and mounted as below, or extensively washed with DMEMF12, L-glu, Pen-Strep antibiotic, and fixed 4h, 8h, 15h, or 24h later.

For ANXA5 experiment, ANXA5 (1mg/ml) was added to the synaptosome suspension (1:20). The mix was added to primary tanycytes for 1h or 4h. Cells were then washed three time with PBS 1X before being fixed with PFA4% for 5 minutes at RT, stained, and mounted as above.

### Sample collection from cell culture

For qPCR, cells were washed twice with ice-cold PBS before adding 350 μL of lysis buffer (RNeasy Mini Kit, #74104, Qiagen). Cells were scraped and collected into an Eppendorf tube for homogenization.

For Western blot, cells were washed twice with ice-cold PBS, before adding 100 µl of lysis buffer (150M NaCl, 1% NP-40, 50mM Tris-HCl, pH 8.0, 0.5% Sodium deoxycholate, 0.1% SDS, Protease inhibitor, PhosphoStop). Cells were scraped and processed for protein quantification by the BCA protein assay kit (23227, Thermo Fisher).

### Molecular biology

For qPCR, the RNeasy Mini kit (#74104 QIAGEN) was used to extract mRNA from cell culture samples. 100ng for cell culture was then reverse-transcribed using the M-MLV reverse transcriptase (M3683, Promega) following the manufacturer’s instructions. cDNAs were diluted (1:10), and qPCR was performed using GoTaq qPCR Master Mix (Promega). The primers (5ʹ-3ʹ) used in this study are available on demand.

For Western Blot, proteins from cell culture were extracted using a lysis buffer (150M NaCl, 1% NP-40, 50mM Tris-HCl, pH 8.0, 0.5% Sodium deoxycholate, 0.1% SDS, Protease inhibitor, PhosphoStop), and quantified using the BCA protein assay kit (23227, Thermo Fisher). 5 µg of proteins were loaded using Laemmli loading buffer in a 15% Acrylamide/Bis Gel (Tris pH8.8 1.5mol/l, acrylamide/bis 15%, SDS 20%, APS 10%, Temed), migrated for 40min at 90V and then 1h at 120V, and then transferred to a PVDF membrane for 1h at 120V. Immunolabelling of the membrane was then performed using primary antibody overnight in 5% BSA 1x TBS-Tween 0.1% at 4°Cand then with secondary antibodies (1:15’000 in 1x TBS 0.1% Tween). The revelation was done using the LiCor Odyssey Fc. The antibodies and dilutions used in this study are reported in Supplementary Table 1.

### Image analysis

For brain image analyses, the entire mediobasal hypothalamus was divided into four subregions on the anteroposterior axis, corresponding to zone 1 (from bregma -1.3 to -1.6mm), zone 2 (from bregma -1.6 to -1.8 mm), zone 3 (from bregma -1.8 to -2.1 mm), and zone 4 (from bregma -2.1 to - 2.5 mm). These subregions have been previously characterized^32^. All quantifications were performed using ImageJ (Fiji, Version 2.0.0-rc-69/1.52p) or IMARIS (version 9.9.1, Bitplane).

The density of synapses in hypothalamic nuclei was assessed with VGAT/Gephyrin and VGLUT2/PSD95 labeling along the anteroposterior and dorsoventral axes. 63x images were processed using “Z-project” and “8-bit conversion”. The number of pre- and post-synaptic puncta and their colocalization (i.e., synapses) was measured using the ImageJ plugin Puncta Analyzer and normalized over the nucleus area delimited by DAPI.

Synapse engulfment by tanycytes *in vivo* was performed on 63x images uploaded on IMARIS (version 9.9.1, Bitplane). After 3D reconstruction of the tdTomato-positive tanycyte, the percentage of the volume occupied by VGLUT2-positive pre-synaptic element was quantified automatically (*Surface, Mask, Statistic functions*).

For cell culture images, VIM-positive tanycytes and synaptosome engulfment were quantified on 40x images. In brief, images were processed using ImageJ (Fiji, Version 2.0.0-rc-69/1.52p): “8-bit conversion”, “Subtract Background”, “Brightness/Contrast”, “Set threshold”, and converted to a mask. The percentage of tanycyte area (delimited by vimentin staining) covered by tdTomato-positive synaptosome was assessed using “Measure”.

For electron microscopy data interpretation, tanycytes were recognized using the ventricular wall (i.e., cell body lining the ventricular wall), the 3D cell morphology (i.e., the presence of a long process), and their ultrastructural characteristics based on previous studies^32^. For tanycyte-synapse interactions using electron microscopy, 3D-reconstructed images from 11 different tanycytes were considered. The proportion of contacts with pre-terminal, post-terminal, symmetric, or asymmetric synapses per tanycyte was calculated among all observed tanycyte-synapse interactions.

### qPCR analyses

Gene expression levels were normalized to actin (*Act*) using the 2−ΔΔCt method and were presented as relative transcript levels.

### Western blot analyses

Densitometric analysis was performed using Image Studio Lite (Version 5.2).

### Single-cell RNAseq data analysis

Publicly available scRNAseq datasets were analyzed as described in their original papers with a few modifications^6,33,67^. Kim *et al.* developmental hypothalamic scRNAseq dataset^6^ was analyzed as described in Lopez-Rodriguez et al^34^. Briefly, the datasets from Kim et al. (GSE132355)^6^ were analyzed and integrated using Seurat 4.1.1. The dataset contains multiple developmental time points (E10-E16, E18, P4, P14, and P45). Cells were filtered to contain at least 200 features, and all datasets were normalized and scaled using scTransform. Following UMAP dimensional reduction (maintaining the first 50 PCA variables), cells were clustered (resolution=1.7) and cell types were identified using known cell marker genes from the original publication. Volcano plots were generated using normalized expression levels per cell population of interest. A dataset subset was generated to explore expression levels according to ependyma developmental trajectories from E11 to P45 (except P8) and integrated and normalized as described in Lopez-Rodriguez et al^34^. The dataset contained exclusively NPCs, ependymal cells, and tanycytes. Feature developmental expression levels were explored using the FeaturePlot() function from Seurat. Finally, the Hypomap dataset^67^ Seurat object was directly retrieved from CellxGene and explored without modifications. Feature plots were generated to observe expression levels across major cell populations.

### Statistical analysis

All values are expressed as means ± SEM. Data were analyzed for statistical significance with Graph Prism 5 software (Version 11.0), using unpaired t-test, one-way ANOVA followed by a Tukey’s post-hoc test, or two-way ANOVA followed by a Bonferroni’s post-hoc test when appropriate. P-values of less than 0.05 were considered to be statistically significant. Statistics are presented in the figures. The statistical tests used, and the numbers of n, cohorts, or cultures are indicated in the legends.

